# A new perspective on HIV cure

**DOI:** 10.1101/007294

**Authors:** Florian Hladik

**Affiliations:** Departments of Obstetrics and Gynecology, and Medicine, University of Washington, and the Vaccine and Infectious Disease Division, Fred Hutchinson Cancer Research Center, Seattle, USA

## Introduction

Current antiretroviral treatment (ART) is highly effective in controlling HIV replication and in many patients suppresses the number of virions measurable in peripheral blood, i.e., the HIV viral load, to undetectable levels. Nevertheless, whenever ART is stopped, HIV levels rebound and the disease returns. This lack of eradication is attributed to a stable latent reservoir of HIV-1 in resting CD4^+^ T lymphocytes and perhaps other susceptible cell types such as macrophages^1^. These cells harbor HIV in the form of proviruses that are integrated into the host chromosomal genome. During effective ART the decay rate of this reservoir is so slow that it would theoretically require treatment for 60 years or longer to eliminate it^2^.

For this reason, HIV-related research efforts and funding are increasingly being devoted to understanding the nature of this latent virus reservoir and how to eradicate it. Two aspects of the latent virus reservoir have emerged as crucial in maintaining infection. First, HIV is not transcribed and translated from some latently infected cells, allowing them to escape detection from the immune system. Second, cells with integrated provirus persist and even expand despite continuous ART^3–5^. To circumvent viral persistence, “kick and kill” strategies have been proposed that attempt to reactivate HIV with latency-reversing agents and then destroy these cells with the help of targeted active or passive immunization strategies. Unfortunately, reactivating cells from infected individuals *ex vivo* has thus far not shown promising results^6, 7^.

In this Perspective, I provide the rationale for my hypothesis that nucleoside reverse transcriptase inhibitors prevent HIV cure by promoting the survival of cells with integrated provirus. If correct, we may be closer to a cure than we realize.

Why CD4^+^ T cells carrying HIV proviruses continue to expand during ART yet viral proteins are not expressed in the process remains an unresolved paradox. To solve this puzzle, tremendous effort is going into characterizing the latent virus reservoir on the one hand and understanding ongoing immune activation during ART on the other. It is hoped that a synthesis of findings in both areas may provide important clues about navigating available and newly arising treatment options toward a cure.

## Mucosal effects of tenofovir

A third area that has been little considered is the effect of ART drugs, both on viral latency and immune activation. Modern antiretroviral combination therapy provides tremendous clinical benefits for HIV-infected patients, dramatically improving quality of life and prolonging life expectancy. Thus, the possibility that a component of ART could paradoxically decrease the chance of cure has never been considered. I was of the same mindset in 2008 when we initiated a systems biology evaluation of the effect of a microbicide gel on the rectal mucosa in a phase I clinical safety trial. This trial, MTN-007, tested the safety and tolerability of a rectal gel formulation containing 1% tenofovir, a phosphonated nucleoside reverse transcriptase inhibitor (NRTI) in development for potential use to prevent rectal HIV transmission^8^. A gel containing 2% nonoxynol-9 (N-9), a temporary mucosal toxin, was included as a positive control arm, and hydroxyethyl cellulose gel and no gel served as negative controls. Our original hypothesis was that the effects of 1% tenofovir gel on the mucosal transcriptome would be negligible whereas N-9 would stimulate inflammatory signatures. However, upon unblinding of the microarray data, we were surprised to find that tenofovir caused many more genes to change than N-9, more often suppressing than enhancing gene expression^9, 10^.

Tenofovir caused three particular changes that bear potential relevance to the HIV cure agenda. First, it strongly inhibited the transcription of a large number of nuclear transcription factors; second, it inhibited the anti-inflammatory function of mucosal epithelial cells; and third, it stimulated signatures of increased cell proliferation and viability. Results obtained from rectal biopsies were replicated in primary vaginal epithelial cells, which also proliferated significantly faster in tenofovir’s presence. In addition to the breadth of transcriptional changes, individual effects caused by tenofovir were not subtle. For example, both in vivo and in vitro, the drug blocked transcription and protein production of interleukin 10 (IL-10) in the range of 90% (Hladik F. et al., submitted for publication).

## An emerging hypothesis

From these data grew my first suspicion that tenofovir, and perhaps more generally NRTIs, could have unappreciated effects on HIV latency, and may in fact prevent HIV cure by promoting the survival of cells with integrated provirus (Figure 1). In developing this concept further below, I want to caution that many of the statements are preliminary and/or hypothetical, intended to serve as a stimulus to the field for further investigation and verification.

**Figure 1.**
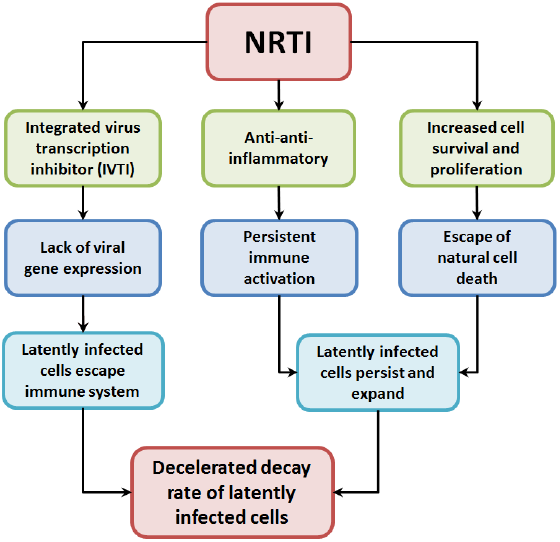
Hypothesized effects of nucleoside reverse transcriptase inhibitors (NRTI) on HIV latency.

Based on the pronounced inhibitory activity of tenofovir on transcription of many genes, I hypothesize that it also inhibits transcription of provirus integrated in such genes. Host gene transcriptional activity has been shown to be an important determinant of integrated HIV transcription^11^. This “integrated virus transcription inhibitor” (IVTI) effect of tenofovir and other NRTIs could explain the transcriptional silence of integrated provirus during ART, since nearly all patients receive an ART regimen containing not just one but two NRTI drugs. Tenofovir’s IVTI activity is circumstantially supported by the following findings. Tenofovir also inhibits herpes simplex virus (HSV) replication^12^, underscoring that its antiviral action *in vivo* is not confined to suppressing the reverse transcriptase of a retrovirus. More intriguingly, genes reported in two recent studies to be preferential sites of HIV integration after periods of ART appear to overlap with genes inhibited in our studies by tenofovir^3, 5^. The list of genes highlighted in the two papers and found to be strongly inhibited by tenofovir in the rectum included CREBBP, IL6ST, KIF1B, FBXW7, DDX6, IKZF3, ZNF652, DST, CLIC5, GRB2, CEPT1, TAOK1 and PAK2. No overlap was found with genes inhibited by N-9. An extended and more formal analysis of these data sets will be interesting. If true, this overlap would imply that over time NRTIs select for cells in which latent HIV survives because of the drugs’ inhibitory effects on transcription of genes hosting integrated provirus.

The IVTI function of NRTIs could be complemented in favoring latency by the drugs’ inhibitory effect on the immune system’s anti-inflammatory circuits. In our study, tenofovir was not directly inflammatory, but its strong inhibition of IL-10, as well as of pathways downstream of the immune homeostatic factor TGF-β, indicated that once inflammation is triggered by an outside event, which could be HIV infection itself, it could be prolonged or perpetuated in the presence of tenofovir. I call this the anti-anti-inflammatory action of tenofovir.

In our cohort of individuals at low risk for inflammation and HIV infection, we did not detect overt inflammation, although tenofovir did significantly increase the density of CD3^+^ and CD7^+^ lymphocytes in the rectal mucosa. In CAPRISA 004, an efficacy trial that demonstrated an overall 39% protective effect of vaginal 1% tenofovir gel^13^, participating women were at much higher risk for inflammation and HIV infection. This uncovered an interesting paradoxical effect of tenofovir: in the presence of inflammation the risk of HIV infection increased significantly more in the tenofovir arm than the placebo arm (personal communication, Dr. Jo-Ann Passmore, University of Cape Town – submitted for publication). This effect was in fact strikingly strong and I hypothetically attribute it to tenofovir’s anti-anti-inflammatory action.

Thus, the anti-anti-inflammatory effect of NRTIs could explain why the massive immune activation caused by primary HIV infection never completely reverses despite effective ART. Interestingly, a similar persistence of immune activation is observed in HSV infection treated with acyclovir, a nucleoside analogue related to NRTIs, which also inhibits DNA synthesis by terminating the growing strand. That HSV-induced local immune activation does not resolve well with acyclovir treatment has been identified as a possible reason why the HSV-associated increase in HIV susceptibility does not reverse when women with genital HSV infection receive acyclovir^14, 15^. Perhaps acyclovir has some of the same anti-anti-inflammatory properties as tenofovir.

Residual immune activation perpetuated by NRTIs could drive the ongoing expansion of cells harboring integrated provirus, and their IVTI function could simultaneously limit transcription of these proviruses. Indeed, our analysis so far indicates that the genes generally turned on by cell activation and the HIV-hosting genes inhibited by tenofovir are different, potentially explaining this apparent paradox. Additionally, the direct cell proliferation- and viability-enhancing effects of tenofovir could contribute to the persistence of latently infected cells.

## ART and anatomic sites of HIV latency

In principle, the HIV latency-inducing effects of NRTIs would likely be strongest where drug concentrations are highest *in vivo*. Studies on tenofovir’s bio-distribution after oral administration show that it highly enriches in gut tissues^16–18^, where the latent HIV reservoir is believed to reside^19^. Estimates also indicate that the rectal concentrations of tenofovir diphosphate, the active intracellular metabolite, are comparable after a single dose of intrarectal 1% tenofovir gel or seven days of oral dosing (personal communication, Dr. Craig Hendrix, Johns Hopkins University). Thus, it is likely that some of the effects we observed in the rectal mucosa after topical application also occur after oral dosing, in particular with years of administration and in combination with a second NRTI.

Of note, after oral dosing, NRTI drug concentrations may be even higher in the small intestine than in the colon and rectum, because in the upper GI tract locally dissolving drug likely adds to drug distributing from the blood stream. If NRTIs do indeed promote latency, then high NRTI concentrations would make the small intestine favorable for HIV latency, consistent with the observation that within the gut the duodenum and ileum were preferential sites of residual HIV DNA and unspliced RNA in ART-suppressed patients^19, 20^. In fact, if NRTIs did not enhance latency, it would be difficult to explain why residual HIV is found precisely where antiretroviral drug concentrations are highest.

## Cure without ART

Circumstantial evidence suggests that ART is not required to cure HIV/SIV infection. The only adult patient ever cured of HIV infection, the “Berlin patient” Timothy Brown, received a stem cell transplant from a donor homozygous for a 32-bp deletion in the CCR5 allele, which provides resistance against HIV-1 infection^21^. He took suppressive ART until the point of his first stem cell transplant, at which point he stopped all ART and never resumed it. Of course, he received a powerful alternative to ART in the form of two CCR5-deficient stem cell transplants, carried out about one year apart. However, he did not achieve complete chimerism for some time after transplantation, because CCR5 receptor-expressing macrophages were still present in rectal biopsies 5.5 months following the stem cell transplants^21, 22^, and thus potential HIV target cells were not completely eliminated at that point. This could have provided a hold for residual HIV. Perhaps removing the hypothetical latency-favoring activity of the NRTI drugs could have contributed to his cure.

In contrast, two HIV-1-infected patients in Boston who also received stem cell transplants continued ART in the peri- and post-transplantation period, and were not cured^23^. While these two patients did not receive CCR5-negative stem cells, which provided a much less favorable scenario than in the Berlin patient’s case, the fact that they continued ART exemplifies the mindset that a novel cure strategy should always be administered in conjunction with the standard of care.

The only animals ever cured from a highly pathogenic SIV infection were rhesus macaques who had been vaccinated before SIV challenge with SIV-protein-expressing rhesus cytomegalovirus vectors^24^ (I exclude animal models and human cases of extremely early ART after infection from the cure definition, since in these cases a latent virus reservoir was likely never established). Although the vaccinated rhesus macaques all showed signs of ongoing systemic infection for weeks or months after challenge, protected monkeys lost all indications of SIV infection over time, consistent with immune-mediated clearance of an established lentivirus infection. None of these animals ever received ART.

While it was suggested that establishment of a latent SIV reservoir might have been prevented by the persistently high frequencies of vaccine-induced SIV-specific CD8^+^ T lymphocytes, early on many of these animals showed clear signs of productive infection, which requires viral integration. Thus, a latent reservoir was likely established, but, hypothetically, in the absence of the latency-prolonging effects of NRTIs the decay rate of provirus-containing cells was accelerated, due to faster natural cell death, less cell expansion, and higher expression of viral proteins, allowing immune recognition by the SIV-specific cytolytic T cells. No viral blips were detected in any animals beyond 70 weeks, perhaps offering a clue as to the time frame required to eradicate a latent reservoir in the absence of NRTIs. However, the pool of latently infected cells was likely small in these animals, and eradication of a larger reservoir may take longer.

## Conclusion and outlook

In summary, given that (1) NRTIs may prevent immune detection of latently infected cells by inhibiting transcription of integrated virus, (2) NRTIs may increase persistence of cells with integrated virus by perpetuating inflammation and enhancing cell proliferation, (3) the only monkeys ever cured of SIV infection never received ART, and (4) the only adult patient ever cured of HIV infection discontinued ART before initiating another powerful antiviral therapy, I hypothesize that effectively suppressing HIV with a strategy that does not contain an NRTI component has curative potential.

Only a few years ago, finding a similarly suppressive alternative to an NRTI-containing ART regimen would have posed a dilemma. Today, powerful second-generation integrase inhibitors and non-NRTI drugs (NNRTIs) are entering early human trials, therapeutic immune strategies using active vaccination and passively infused neutralizing antibodies show promising results, and even more complex strategies such as HIV receptor deletion and specific destruction of integrated viral DNA sequences are progressing. We are thus moving into a phase where effective NRTI-sparing strategies are becoming reality and could offer hope for a cure.

One phase IIb trial, NCT02120352, will soon begin to enroll HIV-1-infected patients who are initially suppressed with an NRTI-containing regimen and then switch to an NRTI-free combination of GSK744 LA, a long-acting injectable formulation of the novel integrase inhibitor GSK1265744^25^ and TMC278 LA, a long-acting injectable formulation of the novel NNRTI TMC278 (ripilvirine)^26^. Though not designed to test a cure, this regimen may, in fact, have curative potential. The study sponsors should consider adjusting their design for that purpose.

## Acknowledgements

I thank M. Juliana McElrath, Fred Hutchinson Cancer Research Center, for helpful discussions, and Sean Hughes, University of Washington, for editing the manuscript.

## References

1 Siliciano RF (2014) Opening Fronts in HIV Vaccine Development: Targeting reservoirs to clear and cure. Nat Med 20(5):480–481.

2 Finzi D, et al. (1999) Latent infection of CD4+ T cells provides a mechanism for lifelong persistence of HIV-1, even in patients on effective combination therapy. Nat Med 5(5):512–517.

3 Maldarelli F, et al. (2014) HIV latency. Specific HIV integration sites are linked to clonal expansion and persistence of infected cells. Science 345 (6193):179–183.

4 Kearney M, et al. (2014) Massive expansion of HIV infected cells with identical proviruses in patients on suppressive ART. 21st Conference on Retroviruses and Opportunistic Infections, p 390.

5 Wagner TA, et al. (2014) Proliferation of cells with HIV integrated into cancer genes contributes to persistent infection. ScienceXpress doi:10.1126/science.125630.

6 Bullen CK, Laird GM, Durand CM, Siliciano JD, & Siliciano RF (2014) New ex vivo approaches distinguish effective and ineffective single agents for reversing HIV-1 latency in vivo. Nat Med 20(4):425–429.

7 Cillo AR, et al. (2014) Quantification of HIV-1 latency reversal in resting CD4+ T cells from patients on suppressive antiretroviral therapy. Proc Natl Acad Sci U S A 111(19):7078–7083.

8 McGowan I, et al. (2013) A phase 1 randomized, double blind, placebo controlled rectal safety and acceptability study of tenofovir 1% gel (MTN-007). PLoS ONE 8(4):e60147.

9 Hladik F, Ballweber L, Dai J, McElrath MJ, & McGowan I (2013) Tenofovir 1% gel causes strong and broad changes to the mucosal transcriptome in a phase I rectal microbicide trial. 16th International Congress of Mucosal Immunology, p T.78.

10 Hladik F, Ballweber L, McElrath MJ, & McGowan I (2013) MTN-007: Integrating systems biology into microbicide trials. Microbicide Trials Network Annual Meeting, Plenary Session Presentation, available from http://www.mtnstopshiv.org/2013annualmeeting.

11 Han Y, et al. (2008) Orientation-dependent regulation of integrated HIV-1 expression by host gene transcriptional readthrough. Cell Host & Microbe 4(2):134–146.

12 Andrei G, et al. (2011) Topical tenofovir, a microbicide effective against HIV, inhibits herpes simplex virus-2 replication. Cell Host & Microbe 10(4):379–389.

13 Abdool Karim Q, et al. (2010) Effectiveness and safety of tenofovir gel, an antiretroviral microbicide, for the prevention of HIV infection in women. Science 329(5996):1168–1174.

14 Celum C, et al. (2008) Effect of aciclovir on HIV-1 acquisition in herpes simplex virus 2 seropositive women and men who have sex with men: a randomised, double-blind, placebo-controlled trial. Lancet 371 (9630):2109–2119.

15 Zhu J, et al. (2009) Persistence of HIV-1 receptor-positive cells after HSV-2 reactivation is a potential mechanism for increased HIV-1 acquisition. Nat Med 15(8):886–892.

16 Patterson KB, et al. (2011) Penetration of tenofovir and emtricitabine in mucosal tissues: implications for prevention of HIV-1 transmission. Sci Transl Med 3(112):112re114.

17 Hendrix CW, et al. (2014) Tenofovir-emtricitabine directly observed dosing: 100% adherence concentrations (HPTN 066). 21st Conference on Retroviruses and Opportunistic Infections, p 104.

18 Louissaint NA, et al. (2013) Single Dose Pharmacokinetics of Oral Tenofovir in Plasma, Peripheral Blood Mononuclear Cells, Colonic and Vaginal Tissue. AIDS Res Hum Retroviruses 29(11):1443–50.

19 Chun TW, et al. (2008) Persistence of HIV in gut-associated lymphoid tissue despite long-term antiretroviral therapy. J Infect Dis 197(5):714–720.

20 Yukl SA, et al. (2010) Effect of raltegravir-containing intensification on HIV burden and T-cell activation in multiple gut sites of HIV-positive adults on suppressive antiretroviral therapy. AIDS 24(16):2451–2460.

21 Hutter G, et al. (2009) Long-term control of HIV by CCR5 Delta32/Delta32 stem-cell transplantation. N Engl J Med 360(7):692–698.

22 Allers K, et al. (2011) Evidence for the cure of HIV infection by CCR5Delta32/Delta32 stem cell transplantation. Blood 117(10):2791–2799.

23 Henrich TJ, et al. (2013) Long-term reduction in peripheral blood HIV type 1 reservoirs following reduced-intensity conditioning allogeneic stem cell transplantation. J Infect Dis 207(11):1694–1702.

24 Hansen SG, et al. (2013) Immune clearance of highly pathogenic SIV infection. Nature 502(7469):100–104.

25 Andrews CD, et al. (2014) Long-acting integrase inhibitor protects macaques from intrarectal simian/human immunodeficiency virus. Science 343(6175):1151–1154.

26 Azijn H, et al. (2010) TMC278, a next-generation nonnucleoside reverse transcriptase inhibitor (NNRTI), active against wild-type and NNRTI-resistant HIV-1. Antimicrob Agents Chemother 54(2):718–727.

